# Labeling of prokaryotic mRNA in live cells using fluorescent *in situ* hybridization of transcript-annealing molecular beacons (FISH-TAMB)

**DOI:** 10.1101/178368

**Authors:** Rachel L. Harris, Maggie C. Y. Lau, Esta van Heerden, Errol Cason, Jan-G Vermeulen, Anjali Taneja, Thomas L. Kieft, Christina DeCoste, Gary Laevsky, Tullis C. Onstott

**Author notes:** Author Correspondence: B80 Guyot Hall, Dept. of Geosciences, Princeton University, Princeton, NJ 08544; (R. L. H.). Author Contributions: M.C.Y.L. is credited for the concept of FISH-TAMB in microbiology. R.L.H, M.C.Y.L., and T.C.O. contributed to experimental design. R.L.H., M.C.Y.L, and A.T. performed *in vitro* hybridization assays. R.L.H., M.C.Y.L, and A.T. were responsible for the maintenance of *E. coli* clone lines. R.L.H. maintained *M. barkeri* and methanogenic enrichment cultures from the BE326 -BH2 borehole. R.L.H. performed *in vivo* hybridization assays, microscopy, and flow cytometry. C.D. assisted R.L.H. in the acquisition of flow cytometry data. G. L. assisted R. L. H. in microscopy. R.L.H., M.C.Y.L., T.C.O., T.L.K., E van H., E.C., and J.V. collected environmental samples from the BE326 -BH2 borehole. All authors contributed in the preparation of this manuscript.

## Abstract

High-throughput sequencing and cellular imaging have expanded our knowledge of microbial diversity and expression of cellular activity. However, it remains challenging to characterize low-abundance, slow-growing microorganisms that play key roles in biogeochemical cycling. With the goal of isolating transcriptionally active cells of these microorganisms from environmental samples, we developed fluorescent *in situ* hybridization of transcript-annealing molecular beacons (FISH-TAMB) to label living prokaryotic cells. FISH-TAMB utilizes polyarginine cell-penetrating peptides to deliver molecular beacons across cell walls and membranes. Target cells are fluorescently labeled via hybridization between molecular beacons and messenger RNA of targeted functional genes. FISH-TAMB’s target specificity and deliverance into both bacterial and archaeal cells were demonstrated by labeling intracellular methyl-coenzyme M reductase A (*mcr*A) transcripts expressed by *Escherichia coli* mcrA^+^, *Methanosarcina barkeri,* and a methanogenic enrichment of deep continental fracture fluid. Growth curve analysis supported sustained cellular viability following FISH-TAMB treatment. Flow cytometry and confocal microscopy detected labeled single cells and single cells in aggregates with unlabeled cells. As FISH-TAMB is amenable to target any functional gene of interest, when coupled with cell sorting, imaging, and sequencing techniques, FISH-TAMB will enable characterization of key uncharacterized rare biosphere microorganisms and of the syntrophically activated metabolic pathways between physically associated microorganisms.

## INTRODUCTION

The studies of ribosomal RNA (rRNA), whole-genome and shotgun sequencing have vastly expanded our knowledge of microbial diversity and metabolic potential in natural communities. Recently, sequencing technologies have unveiled genomic and functional information on uncultivable microbial species, including rare biosphere taxa and microbial dark matter (MDM) (Sogin *et al.*, 2006; Rinke *et al.*, 2013; Spang *et al.*, 2015; Seitz *et al.*, 2016; Vanwonterghem *et al.*, 2016; Lazar *et al.*, 2017; Zaremba-Niedzwiedzka *et al.*, 2017). The metabolic potential and versatility encoded in genomes are useful for predicting organisms’ adaptability to shifts in environmental conditions. However, it remains challenging to determine actual roles in biogeochemical cycling under *in situ* conditions. Metatranscriptomic analysis does reveal *in situ* metabolic activity of microbial ecosystems. However, the analysis of unlinked mRNA sequences in metatranscriptomic data sets renders it difficult to accurately infer taxonomy from functional genes, especially for organisms acquiring functional genes via horizontal gene transfer (HGT). The taxonomic identity of single cells from environmental samples is typically performed through fluorescence *in situ* hybridization (FISH), which involves the use of fluorescent oligonucleotide linear probes targeting the 16S rRNA gene (DeLong *et al.*, 1989; Amann *et al.*, 1990). Because ribosomes are far more abundant in metabolically active versus inactive microorganisms, FISH also determines the abundance of active microbial constituents relative to the total community (Karner and Fuhrman, 1997; Williams *et al.*, 1998; Christensen *et al.*, 1999; Pernthaler *et al.*, 2002). However, 16S rRNA-based identification requires prior knowledge of the target organisms’ phylogenies and provides no direct evidence of the target organisms’ metabolic roles.

FISH methods targeting messenger RNA (mRNA) in prokaryotes have been developed to relate functional gene expression within organisms to their metabolic contributions to biogeochemical cycling (Pernthaler and Amann, 2004; Kalyuzhnaya *et al.*, 2006; Jen *et al.*, 2007; Mota *et al.*, 2012). However, these studies were performed on fixed (i.e. dead) cells, offering only a snapshot of past activity in labeled cells based solely on the single target gene being expressed. *In situ*, real-time metabolic activity imaging has only been applied to genetically engineered strains expressing reporter proteins such as green-fluorescence proteins (Golding *et al.*, 2005). Imaging and sorting of translationally active cells from environmental samples has recently been achieved through the use of Bioorthogonal noncanonical amino acid tagging (BONCAT) (Hatzenpichler *et al.*, 2016). When this staining technique is combined with standard FISH methods, in principle, the taxonomic identity of all active microbial cells in environmental samples can be determined. Since microorganisms that exert significant impacts on their environments include slow-growing and low-abundance taxa, to fully understand their metabolic requirements and hence their function in biogeochemical cycling, it is necessary to strategically select for these key players and study their overall expression profiles. It would be advantageous to use a method that targets metabolically active cells that exhibit certain specific functions, and meanwhile maintains their viability. To our knowledge such a method has not yet been reported.

We describe here the development of fluorescent *in situ* hybridization of transcript-annealing molecular beacons (FISH-TAMB) to target mRNA in viable and transcriptionally active prokaryotic cells. Molecular beacons (MBs), with a hairpin oligonucleotide sequence outfitted with a fluorophore and a fluorescence quencher (Tzschaschel *et al.*, 1996), were selected to target the mRNA of Bacteria and Archaea, because they result in a higher signal-to-background noise ratio than linear probes and have been successfully applied to detect intracellular mRNA of living eukaryotic cells (Sokol *et al.*, 1998; Nitin *et al.*, 2004; Santangelo *et al.*, 2006; Bao *et al.*, 2009; Larsson *et al.*, 2012). In the unbound state, complementary bases on the 5’ and 3’ ends of MBs self-anneal to form a stem structure, which results in fluorescence quenching. Recognition of a target sequence results in MB linearization for subsequent hybridization (Fig. 1). Thus, the fluorophore is no longer in physical proximity to the quencher, resulting in emission of a known wavelength at a level differentiable from the background fluorescence due to autofluorescence and unbound MBs (Goel *et al.*, 2005). In order to deliver the MBs into prokaryotic cells without causing cell lysis, cell-penetrating peptides (CPPs) are used as the cargo-delivering vehicle, as they have been shown to successfully non-lethally deliver DNA and nanoparticles into living cyanobacteria (Liu et al. 2013a,b).

**Fig. 1.**
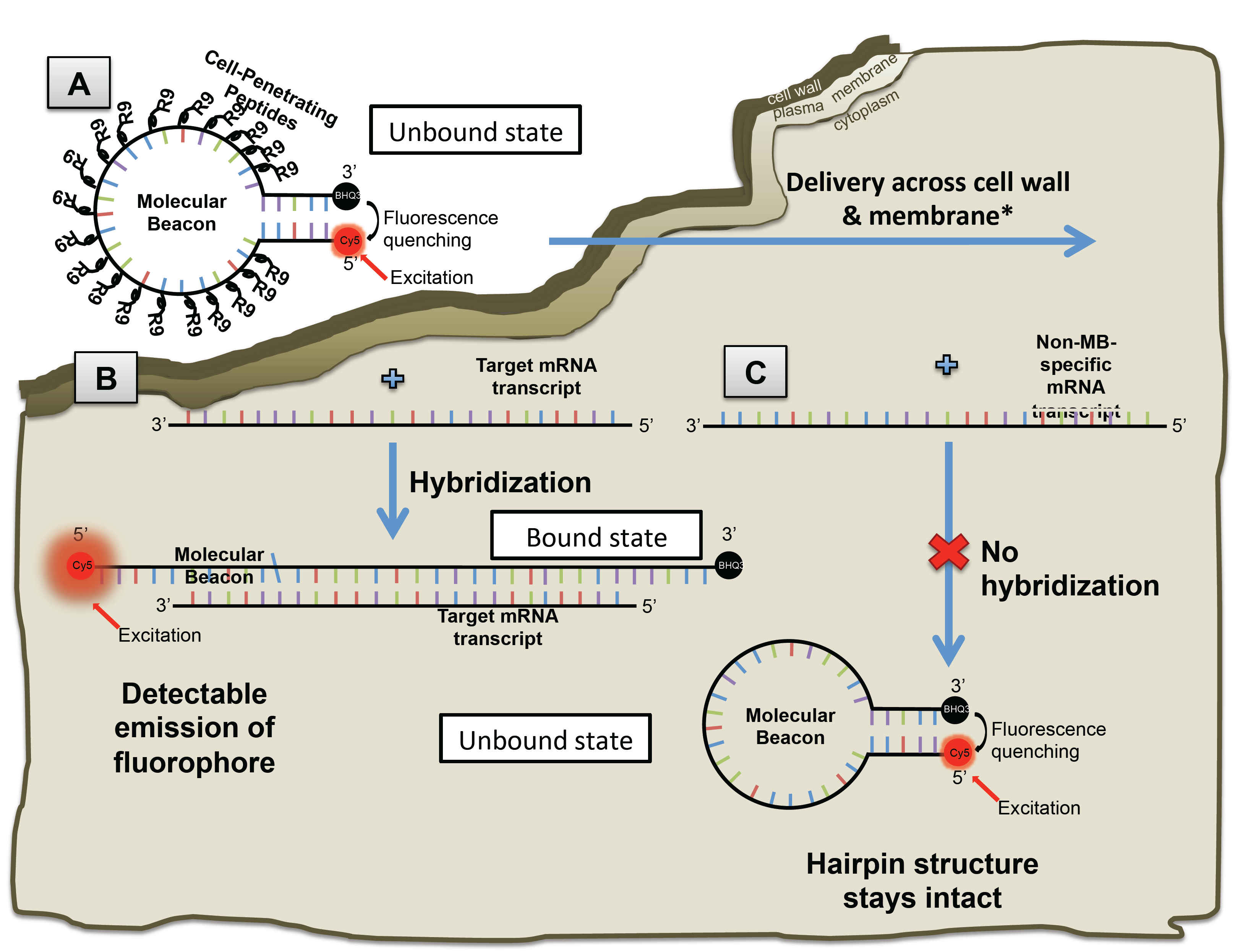
FISH-TAMB probe conformation and hybridization to encountered messenger RNAs. **(A)** An oligomer comprised of a 24 base-long complementary *mcr*A mRNA sequence is flanked by 5 reverse complement nucleotides to form a molecular beacon (MB) loop and stem structure. Cell-penetrating peptides (CPPs) comprising 9 arginine sequences (R9) are non-covalently bound to the MB sequence and are responsible for its delivery across the cell wall and plasma membrane. **(B)** Fluorescence of Cy5 fluorophore covalently bound to the 5’ end of the MB sequence remains quenched by BHQ3 bound to the 3’ terminus until the MB hybridizes to a target transcript sequence. Hybridization results in the linearization of the MB, subsequently unquenching Cy5 from BHQ3, allowing the fluorophore’s emission upon excitation by a source in the red bandwidth of the visible light spectrum. **(C)** If the MB encounters an mRNA transcript that is not its intended target, it will retain its hairpin conformation and fluorescence of Cy5 will remain quenched by BHQ3. Images not to scale. **\***Mechanism of CPP delivery across the cell wall and plasma membrane remains under debate. Intracellular fate of R9 is unknown.

In this study, we employed MBs targeting the gene encoding the alpha subunit of methyl-coenzyme M reductase (*mcr*A), a marker gene of methanogens (Lueders *et al.*, 2001; Luton *et al.*, 2002; Evans *et al.*, 2015; Vanwonterghem *et al.*, 2016) and the uncultivated anaerobic methanotrophs (ANMEs) (Hallam *et al.*, 2003). We examined this FISH-TAMB method in three phases by (i) performing *in vitro* experiments to investigate the fluorescence strength of the *mcr*A FISH-TAMB probe in the absence and presence of *mcr*A mRNA oligonucleotides under different buffer conditions; (ii) performing *in vivo* experiments to validate the delivery of FISH-TAMB probes into living prokaryotic cells; and (iii) evaluating the efficiency of labeling *Escherichia coli* mcrA^+^ expression clones, *Methanosarcina barkeri*, and methanogenic cells enriched from Precambrian shield fracture fluid collected from 1.34 km below land surface (BE326 BH2 borehole) with the *mcr*A FISH-TAMB probes. The effect of FISH-TAMB treatment on cell viability was assessed via growth curve analysis.

## MATERIALS AND METHODS

### Molecular beacon probe design

The MB utilized in this study comprised a GC-rich 5-base pair stem and a 24-mer nucleotide probe sequence (5’-[Cy5]cctggCGTTCATBGCGTAGTTVGGRTAGTccagg[BHQ3]-3’) modified from the mcrA-rev reverse primer (5’-CGTTCATBGCGTAGTTVGGRTAGT-3’) commonly used in diversity studies of methanogens (Steinberg and Regan, 2008). As the additional bases on the stem structure (as indicated by small letters in the MB probe sequence) may affect the specificity of the MB probe to target *mcr*A genes, the similarity between MB probe sequence and *mcr*A genes was verified *in silico* using BLAST (https://blast.ncbi.nlm.nih.gov/Blast.cgi) against the nucleotide database. The MB probe sequence was flanked by a Cy5 fluorophore (excitation peak at 640 nm, and emission peak at 665 nm) covalently bound to the 5’ end and a BHQ3 Black Hole Quencher^^®^^ on the 3’ end (MilliporeSigma, St. Louis, MO USA).

### Formation of R9:MB complexes (FISH-TAMB probes)

A CPP comprised of nine arginine amino acid residues (R9) was selected as a carrier to deliver the MB across cell walls and plasma membranes, as it has been demonstrated to penetrate cyanobacterial walls and membranes without harmful effects (Liu et al. 2013). R9 was mixed with MB in 1x Dulbecco’s phosphate buffered saline solution (DPBS) (Corning Mediatech, Manassas, VA USA) to achieve the following molar ratios: 0:1, 5:1, 10:1, 15:1, 20:1, 25:1, 30:1. Reactions were incubated for 30 minutes at 37°C in a C1000 Touch™ Thermal Cycler (Bio-Rad Laboratories, Inc., Irvine, CA USA). Gel electrophoresis determined the minimum molar ratio of R9 to MB required for complete complexation of all free-floating MB in solution (*SI* Methods).

#### In vitro *hybridization assays*

*In vitro* hybridization assays were performed to assess (i) hybridization of MB and FISH-TAMB probes to target *mcr*A oligonucleotide sequences, (ii) whether resulting fluorescence from probe-target hybridization was differentiable from background fluorescence of unbound and potentially non-specifically bound MB probes, and (iii) optimal incubation time for detection of positive hybridization fluorescence in subsequent experiments. Triplicate 100-μl reaction mixtures containing either 0.4 μM MB or 1 μM FISH-TAMB probes in 1x DPBS were incubated at 37°C for 10 minutes with 0.4 μM of *mcr*A oligonucleotide sequences (5’-ACTAYCCBAACTACGCVATGAACG-3’) complementary to the MB probe sequence. Background signal (due to unbound MB probes) and potential non-specific hybridization fluorescence were respectively assessed by incubating MB and FISH-TAMB probes in the absence of any *mcr*A target (blank) and an oligonucleotide sequence specific to particulate methane monooxygenase beta subunit (*pmo*A) (5’-GAAYSCNGARAAGAACGM-3’) (Luesken *et al.*, 2011). Fluorescence images were taken every 5 minutes for 100 minutes using a Typhoon 9410 Variable Mode Imager^^®^^ (Molecular Dynamics, GE Healthcare, Little Chalfront, UK) (excitation 633 nm, detection bandwidth 655 - 685 nm, exposure time 5 min.).

Fluorescence intensity was measured as a function of temperature and salt concentration to determine stability profiles of the MB probe sequence in the presence and absence of *mcr*A targets (bound MB vs. unbound MB states). Three 50-μl reaction volumes were prepared for the unbound MB controls, comprising 16 nM MB probes and Takara PCR buffer containing 1.5 mM MgCl_2_ (1x), 7.5 mM MgCl_2_ (5x), or 15 mM MgCl_2_ (10x). Three bound MB reactions were set up using the same recipes except with the addition of 32 nM *mcr*A target sequences. Reaction mixtures were incubated at 37°C for 1 hour on a real-time qPCR 7900HT system (Applied Biosystems, Inc., Carlsbad, CA USA). Melting curve analysis was done for temperatures ranging from 25°C to 95°C with fluorescence signals measured every 0.2°C. Optimal detection temperature for positive MB-target hybridization was determined as the temperature with the highest signal-to-background noise ratio, as indicated by the relative fluorescence intensities of bound MB and unbound MB, respectively.

### Microbial sampling of fracture fluid

Fracture fluid was collected in June 2016 following established sampling procedures (Magnabosco *et al.*, 2014; Lau *et al.*, 2014) from a horizontal borehole located 1.34 km below land surface on the 26^th^ level of shaft 3 of the Beatrix Gold Mine in South Africa (BE326 BH2) (S 28.235º, E 26.795º). Due to low *in situ* cell concentration of 10^3^ to 10^4^ cells ml^-1^ (Simkus *et al.*, 2016), the fracture fluid was first filtered using a 0.2 μm hollow fiber MediaKap^^®^^-10 filter (Spectrum Labs, New Brunswick, NJ USA) and then back-flushed with fracture fluid into sterile, N_2_-sparged 160-ml borosilicate serum vials to obtain a final concentration of ∼10^7^ cells ml^-1^. Dissolved gas samples were collected along with field measurements of certain environmental parameters (*SI* Methods, results in Table S3).

#### *Methanogenic enrichments of fracture fluid and* M. barkeri

The methanogenic medium DSMZ medium 120a modified after the recipe of Bryant and Boone (1987) (*SI* Methods) was used to enrich methanogens from the fracture fluid sample with concentrated biomass (BE326 BH2-Conc). The pH of the medium was adjusted to 8.2, the *in situ* pH of the fracture fluid, with anaerobic NaOH prior to inoculation with BE326 BH2-Conc. The DSMZ medium 120a was adjusted to pH 7.2 and inoculated with axenic cultures of methanogen *M. barkeri* (ATCC^^®^^ 43569™). Cultures were incubated at 37°C in the Coy glove bag. Active methanogenesis was verified by detecting novel CH_4_ production in headspace gas using a flame-ionizing detector (FID) gas chromatograph (Peak Performer 1 series, Peak Laboratories, Mountain View, CA USA). Details of methanogenic enrichment maintenance can be found in *SI* Methods.

#### E. coli *expression clones*

The *mcr*A gene was PCR-amplified directly from the *M. barkeri* culture to prepare *E. coli* mcrA^+^ expression clones to serve as a proxy for cells gaining *mcr*A via HGT and to validate the ability of CPP to deliver MBs across bacterial membranes. *E. coli* cells containing a *pmo*A insert (*E. coli* pmoA^+^) isolated from BE326 BH2 fracture fluid served as a negative control for assessing fluorescence due to potentially non-specific FISH-TAMB hybridization *in vivo*. Details regarding the isolation and transformation of *mcr*A and *pmo*A gene inserts into JM109 competent *E. coli* are described in *SI* Methods.

*E. coli* expression clones were monitored for gene loss by periodically plating liquid culture aliquots onto LB/AIX cloning plates for blue/white screening (*SI* Methods). If *mcr*A or *pmo*A was absent from the plasmid, the cloning procedure was repeated prior to FISH-TAMB treatment.

### FISH-TAMB delivery into live prokaryotic cells

All cultures (∼10^6^ cells) including *E. coli* pmoA^+^ were incubated in the dark at 37°C in 100-μl reactions containing 1 μM FISH-TAMB probes in 1x DPBS solution. Reactions for *M. barkeri* and the BE326 BH2-Conc methanogenic enrichment were prepared anaerobically using degassed 1x DPBS in the Coy glove bag to maintain cell activity. Reaction mixtures were placed into a 96 well optical plate (Cellvis, Mountain View, CA USA) and fluorescence emission levels were imaged every five minutes for 120 minutes using the Typhoon 9410 Variable Mode Imager^^®^^ as described above. To assess fluorescence due to potential non-specific MB hybridization *in vivo*, FISH-TAMB probes were also incubated with *E. coli* pmoA^+^.

Growth curve analysis was performed to assess the effect of FISH-TAMB on sustained viability of treated cells relative to untreated cells, with the latter serving as controls. *E. coli* mcrA^+^, *E. coli* pmoA^+^, and *M. barkeri* (∼10^6^ cells) were incubated with 1 μM FISH-TAMB probes as described above and subsequently inoculated into Luria broth containing 0.05 mg ml^-1^ ampicillin (LB/A) and DSMZ 120a media for *E. coli* and methanogenic cultures, respectively. Growth curves were obtained by measuring optical density at 600 nm for *E. coli* using a Beckman DU^^®^^ 530 Life Science UV/Vis Spectrophotometer (Beckman Coulter^^®^^, Indianapolis, IN USA) and at 550 nm for *M. barkeri* using a Hach DR/2010 Spectrophotometer (Hach Company, Loveland, CO USA).

### Enumeration of FISH-TAMB labeled cells by flow cytometry

*E. coli* mcrA^+^, *E. coli* pmoA^+^, *M. barkeri* and BE326 BH2-Conc methanogenic cultures were incubated for 15 minutes at 37°C in 100-μl reaction mixtures containing 1 μM FISH-TAMB probes, 1x DPBS solution and ∼10^6^ cells. *M. barkeri* and BE326 BH2-Conc enrichments were incubated with FISH-TAMB probes anaerobically as described above. Following incubation, the 100-μl reaction mixtures were diluted in 0.9 ml 1x DPBS solution containing ∼10^6^ ml^-1^ Fluoresbrite™ plain red 0.5 μm microspheres (Polysciences, Inc., Warrington, PA USA) for flow cytometric analysis. Flow cytometry was performed on a BD LSRII Multi-Laser Analyzer (Becton, Dickinson and Company, Franklin Lakes, NJ USA) at the Princeton University Flow Cytometry Core Facility. Data were acquired for 120 seconds for each sample at 8 μl min^-1^ average flow rate using four independent laser channels at default wattage settings: 355 nm at 20 mW, 405 nm at 25 mW, 488 nm at 20 mW, and 633 nm at 17 mW. Forward and side-scattered light were set to logarithmic gain and used to trigger events. The system was flushed with 10% (v/v) bleach solution for 1 minute before analysis and between samples of different cell types and after samples treated with FISH-TAMB probes.

Fluorescent microsphere counts were used to calculate the volume of fluids being analyzed and thereby the cell concentrations. For all samples, events gated as cell-sized objects and FISH-TAMB-labeled cells in 1x DPBS + FISH-TAMB probes + growth medium (see *SI* Methods for information regarding cell population gating parameters) were subtracted from final counts collected for each cell type. Statistical analysis of observed differences in FISH-TAMB labeling between samples and their respective controls was performed using a Student t-test (StatPlus:mac LE software, AnalystSoft, Inc., Walnut, CA USA).

### Confocal microscopy

Live-cell imaging was performed on all cell types to qualitatively confirm the ratio of active versus inactive cells enumerated by FISH-TAMB probes in flow cytometry. All cell types were incubated with FISH-TAMB probes as described above. FISH-TAMB-labeled cells were imaged at the Princeton University Confocal Imaging Core using a Nikon Ti-E inverted confocal microscope equipped with a 100X Plan Apo NA 1.45 oil objective lens, Yokogawa CSU-21 spinning disk, and Orca Flash camera (Nikon Instruments, Melville, NY USA). F420 autofluorescence of *M. barkeri* and methanogenic BE326 BH2-Conc cells were excited with the 405-nm laser channel, and detected on the 461-nm emission filter. *E. coli* autofluorescence was excited at 488 nm, and emission was set to 518 nm. Excitation and emission of the Cy5 fluorophore in MB probes were set to 647 nm and 670 nm, respectively. Samples were maintained under a 100% CO_2_ atmosphere during imaging.

A time series imaging experiment was performed to assess Cy5 fluorescence lifetime of FISH-TAMB hybridized cells. Methanogenic BE326 BH2-Conc cells were treated with FISH-TAMB probes as previously described and were imaged every minute for 14 hours using a Nikon Ti-E inverted confocal microscope outfitted with the same equipment as described above with an Andor Zyla sCMOS camera. Imaging parameters were consistent with the previous live-cell imaging experiment.

## RESULTS AND DISCUSSION

### Conformational stability and target specificity

Melting curve analysis revealed maximum fluorescence of bound MB at 25°C under all investigated saline buffer solutions. (*SI* Table S1, Fig. S1). This temperature corresponded with the highest signal-to-background noise ratio (195:1 relative to unbound MB in 1x PCR buffer containing 1.5 mM MgCl_2_). All cell types were maintained at growth temperature (37°C) for 15-minute incubation with FISH-TAMB probes given 133x greater fluorescence of bound MB at this temperature (*SI* Fig. S1) and high fluorescence recorded at this time from an *in vitro* hybridization time series experiment (*SI* Fig. S2). Bound MB fluorescence intensity remained > 17x greater than unbound MB up to 64°C before dropping down to 2x greater emission for higher temperatures up to 95°C (*SI* Fig. S1). MB conformation remained intact at all assessed salinities, but signal-to-background noise improved with increased salt concentration between 55° - 65°C (Table S1). Thus, FISH-TAMB demonstrates a large operational temperature range of 25°C - 65°C, but may be limited from *in situ* studies of thermophiles.

*In vitro* hybridization revealed background autofluorescence of unbound MB significantly diminished when MB was non-covalently bound to R9 (Fig. 2D) and results from gel electrophoresis showed a minimum of 20:1 R9:MB molar ratio for complete complexation of all free-floating MB in solution (*SI* Fig. S3). It is possible that R9 may be playing a role in stabilizing the MB hairpin conformation, thus improving the quencher’s absorption of background fluorophore emission. However, this stabilization appeared to be inhibitory to MB-target hybridization when FISH-TAMB probes were incubated with *mcr*A oligonucleotide sequences *in vitro* (Fig. 2E). Because positive fluorescence signals were detected when FISH-TAMB probes encountered intracellular *mcr*A mRNA *in vivo* after entering cells (Fig. 2F-H), we hypothesize that intracellular scavenging may physically dissociate R9 from MB allowing subsequent MB-target hybridization. While the exact mechanism remains unknown, the MB probes released from R9 appear to retain hairpin conformation following cellular penetration, as evidenced by minimal fluorescence in *E. coli* pmoA^+^ cells incubated with FISH-TAMB probes (Fig. 2I).

**Fig. 2.**
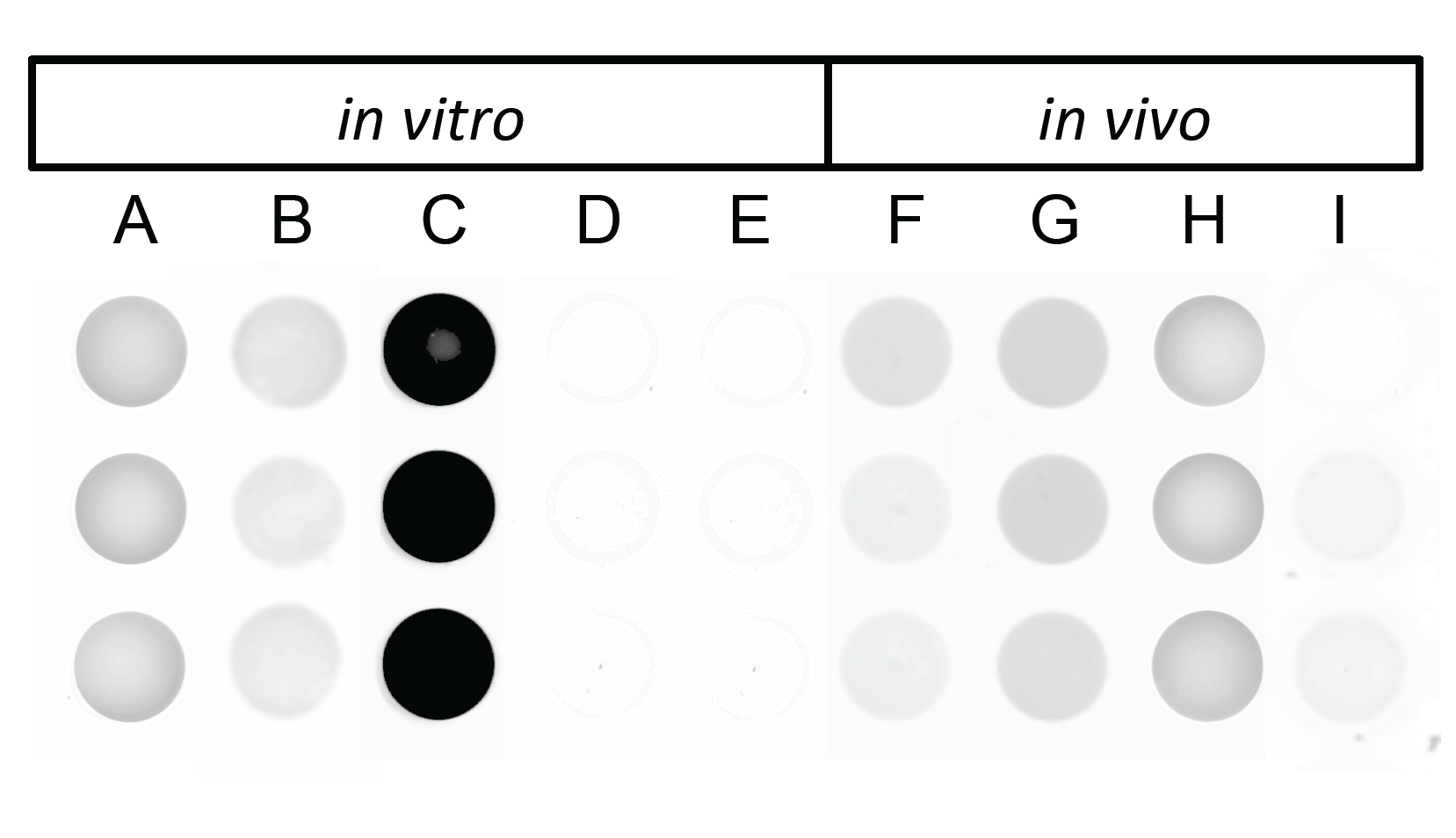
MB and FISH-TAMB probe target specificity *in vitro* and *in vivo*. **(A)** 0.4 μM MB in 1x PBS. **(B)** 0.4 μM MB + *pmo*A target oligo. **(C)** 0.4 μM MB + *mcr*A target oligo. **(D)** 1 μM R9:MB in 1x PBS. **(E)** 1 μM R9:MB + *mcr*A target oligo. **(F)** 1 μM R9:MB *+ M. barkeri*. **(G)** 1 μM R9:MB + BE326 BH2-Conc. **(H)** 1 μM R9:MB + *E. coli* mcrA^+^. **(I)** 1 μM R9:MB + *E. coli* pmoA^+^. Images taken after 20 minutes incubation with a Typhoon 9410 Variable Mode Imager^^®^^. Excitation 633 nm. Emission 675/10 nm. Exposure time 5 minutes.

### Detection of living cells labeled by FISH-TAMB probes

Flow cytometry data revealed that FISH-TAMB-treated cells containing the *mcr*A gene exhibited a significant increase in fluorescence on the Cy5 filter relative to the untreated and *E. coli* pmoA^+^ cell populations (Student t-test, t ≥ 2.6, p < 0.05) (Fig. 3 D-F vs. A-C, Fig. 4, *SI* Table S2). Cells expressing the *mcr*A gene demonstrated a statistically significant difference in the number of FISH-TAMB labeled cells relative to the media blank controls (Student t-test, t ≥ 5.2, p < 0.05) (Table 1). FISH-TAMB-labeled cells of *E. coli* mcrA^+^, *M. barkeri*, and BE326 BH2-Conc represented 2%, 32%, and 1% of the total cell concentration, respectively. The *E. coli* pmoA^+^ negative control exhibited only 0.02% recovery, which was not statistically different from the 1x DPBS + FISH-TAMB probes + growth medium blank (Student t-test, t = 1.6, p = 0.25).

**Table 1.**
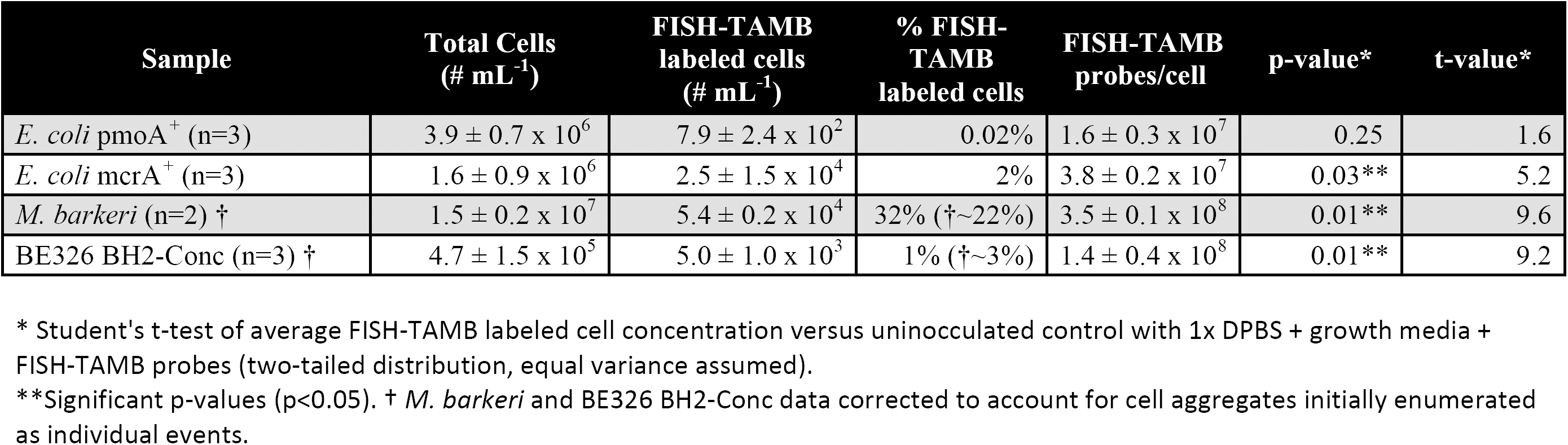
Flow cytometry data of FISH-TAMB treated cells.

| Sample | Total Cells<br>(# mL<sup>-1</sup>) | FISH-TAMB<br>labeled cells<br>(# mL<sup>-1</sup>) | % FISH-TAMB<br>labeled cells | FISH-TAMB<br>probes/cell | p-value* | t-value* |
| --- | --- | --- | --- | --- | --- | --- |
| *E. coli* pmoA<sup>+</sup> (n=3) | 3.9 ± 0.7 x 10<sup>6</sup> | 7.9 ± 2.4 x 10<sup>2</sup> | 0.02% | 1.6 ± 0.3 x 10<sup>7</sup> | 0.25 | 1.6 |
| *E. coli* mcrA<sup>+</sup> (n=3) | 1.6 ± 0.9 x 10<sup>6</sup> | 2.5 ± 1.5 x 10<sup>4</sup> | 2% | 3.8 ± 0.2 x 10<sup>7</sup> | 0.03** | 5.2 |
| *M. barkeri* (n=2) † | 1.5 ± 0.2 x 10<sup>7</sup> | 5.4 ± 0.2 x 10<sup>4</sup> | 32% (†~22%) | 3.5 ± 0.1 x 10<sup>8</sup> | 0.01** | 9.6 |
| BE326 BH2-Conc (n=3) † | 4.7 ± 1.5 x 10<sup>5</sup> | 5.0 ± 1.0 x 10<sup>3</sup> | 1% (†~3%) | 1.4 ± 0.4 x 10<sup>8</sup> | 0.01** | 9.2 |

\* Student's t-test of average FISH-TAMB labeled cell concentration versus uninoculated control with 1x DPBS + growth media + FISH-TAMB probes (two-tailed distribution, equal variance assumed).
\*\*Significant p-values (p<0.05). † *M. barkeri* and BE326 BH2-Conc data corrected to account for cell aggregates initially enumerated as individual events.

**Fig. 3.**
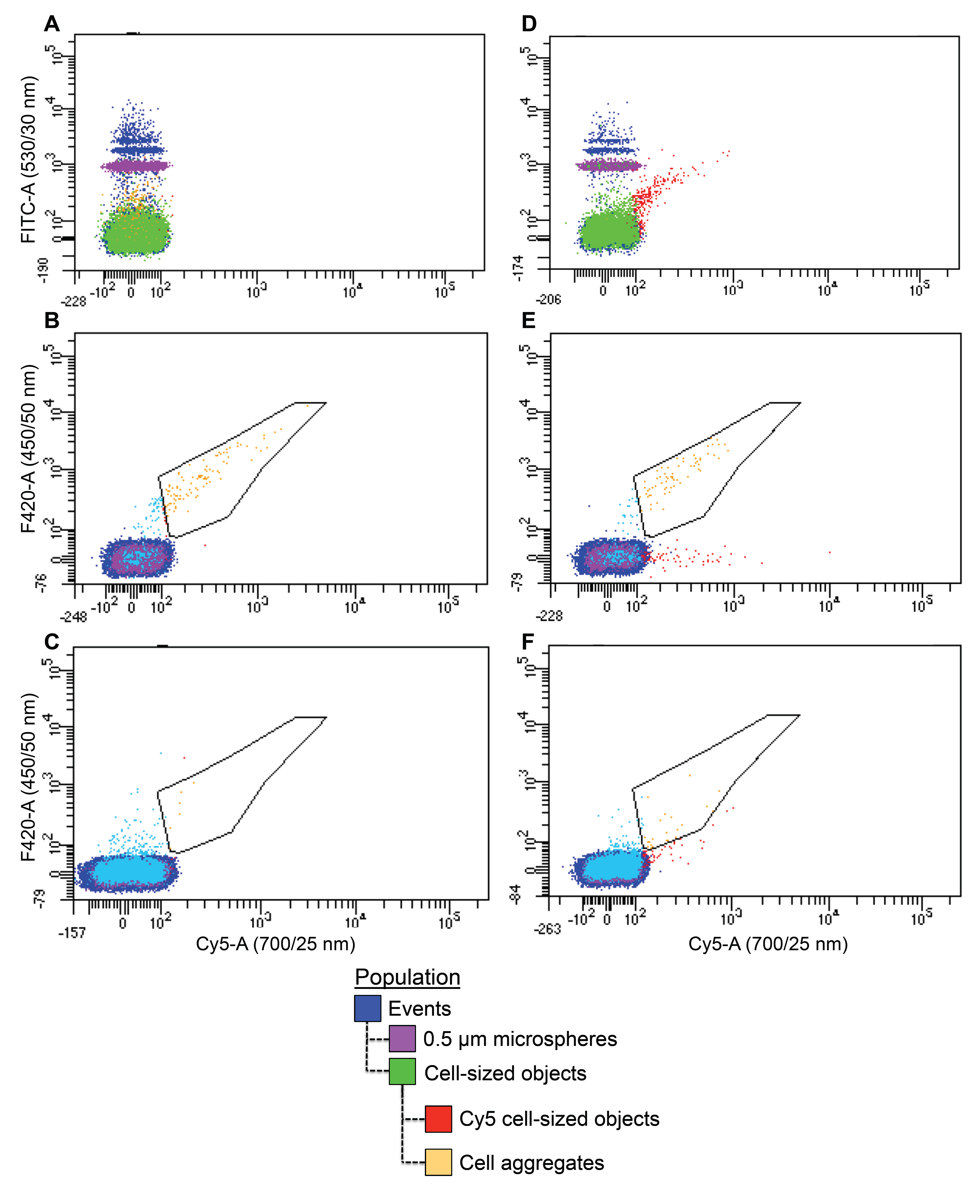
Flow cytometry distinguishes FISH-TAMB labeled cells. Flow cytometry reveals single cells demonstrating distinctive Cy5 fluorescence patterns in cells treated with 1 μM FISH-TAMB probes **(D-F)** relative to untreated cells **(A-C). (A, D)** *E. coli* mcrA^+^. **(B, E)** *M. barkeri.* **(C, F)** BE326 BH2-Conc. Positively labeled cells indicated in red. FISH-TAMB-labeled cells are gated with respect to Cy5 fluorescence intensity (X-axis) and target cell autofluorescence properties (Y-axis).

**Fig. 4.**
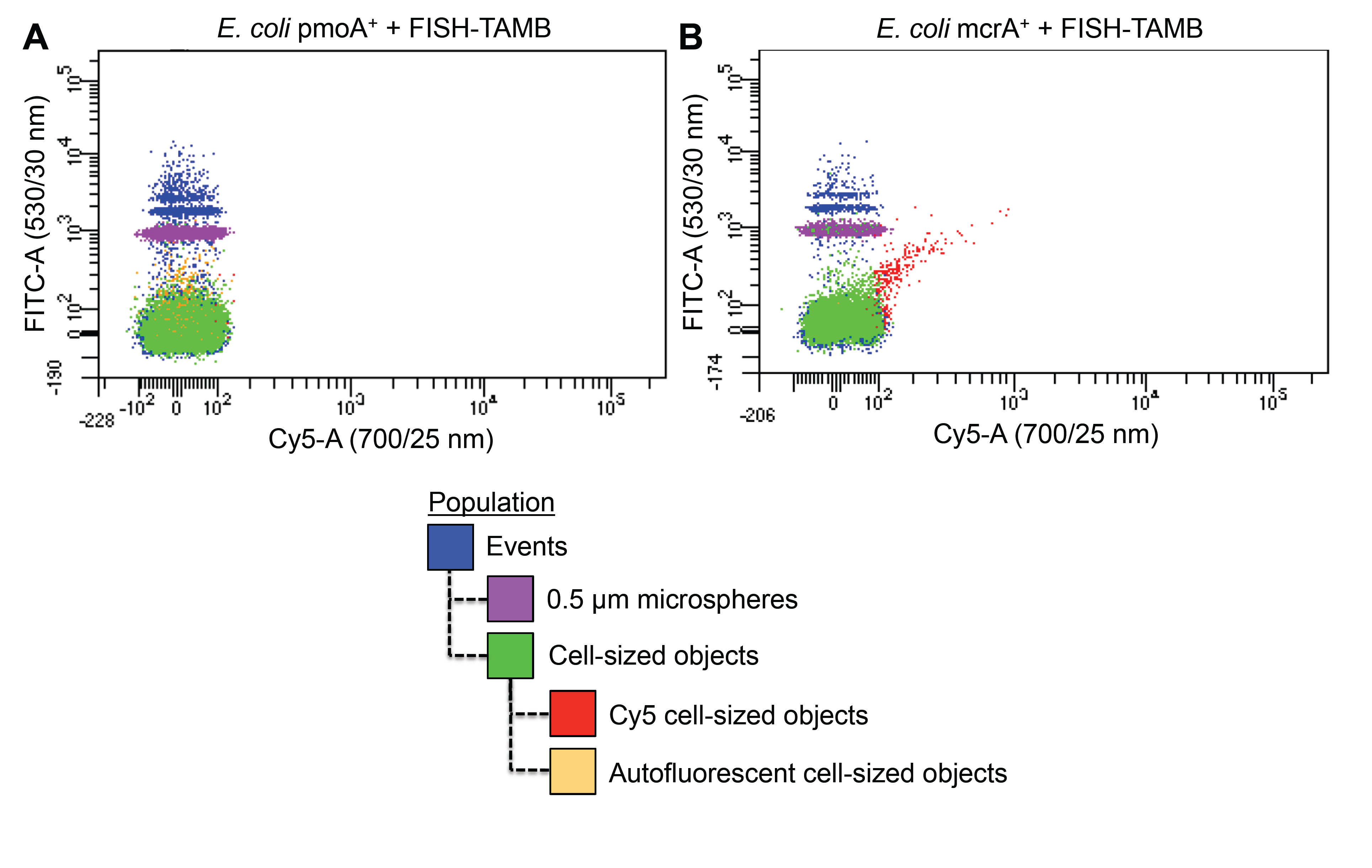
FISH-TAMB specificity to *mcr*A in *E. coli*. **(A)** *E. coli* pmoA^+^ cells lacking the *mcr*A gene do not demonstrate Cy5 fluorescence when treated with 1 μM FISH-TAMB probes. **(B)** Active *E. coli* mcrA^+^ cells demonstrate Cy5 fluorescence when treated with 1 μM FISH-TAMB probes. FISH-TAMB-labeled cells are indicated in red. FISH-TAMB-labeled cells are gated with respect to Cy5 fluorescence intensity (X-axis) and *E. coli* autofluorescence properties (Y-axis).

It was expected that 100% of *E. coli* mcrA^+^ cells would be actively transcribing *mcr*A but only 2% of the cell population was labeled by FISH-TAMB probes. This low recovery of active cells was visually confirmed by spinning disk photomicroscopy, wherein 11 of approximately 600 cells fluoresced under the microscope. Replating non-FISH-TAMB-treated *E. coli* mcrA^+^ cells yielded white colonies with blue centers, suggesting toxicity of the *mcr*A gene insert and its subsequent excision from the plasmid. To evaluate whether promoting *mcr*A expression in *E. coli* would improve the percentage of FISH-TAMB labeled cells, pGEM vectors containing *mcr*A genes were transformed into two alternative overexpression *E. coli* strains, C41(DE3) and BL21(DE3) (*SI* Methods), that are known to be tolerable of unstable and toxic gene inserts (Saïda et al. 2006). However, Typhoon imaging of FISH-TAMB-treated C41(DE3) mcrA^+^ and BL21(DE3) mcrA^+^ cells showed fluorescence emission as low as the C41(DE3) pmoA^+^ cells (data not shown). The reasons for the low percentage of labeled *E. coli* mcrA^+^ observed remain unclear.

In addition to single cells, FISH-TAMB identified active cells in aggregates of *M. barkeri* (Fig. 5A) and BE326 BH2-Conc (Fig. 5E). Cell aggregates were enumerated as individual events (i.e. presumably single cells) by flow cytometry detectors, but were discernable from true single cells by their significantly different FSC-A (Student t-test, t = 52, p < 0.001) and SSC-A (Student t-test, t = 62, p < 0.001) values. FISH-TAMB-labeled cells within aggregates also displayed significantly increased Cy5 fluorescence relative to unlabeled aggregates (Student t-test, t = 12, p< 0.001) (Fig. 3F). Spinning disk photomicroscopy of aggregates revealed that FISH-TAMB labeled ∼20% of cells in *M. barkeri* and ∼46% in BE326 BH2-Conc. Thus, we corrected FISH-TAMB labeling to ∼22% from 32% in *M. barkeri* and ∼3% from 1% in BE326 BH2-Conc (Table 1). Interestingly, the majority of FISH-TAMB labeled cells occurred only within the interior of the *M. barkeri* aggregate. Given this distribution of labeled cells and an average incubation ratio of 1.4 ± 0.1 x 10^8^ probes for every cell (Table 1), it is unlikely that FISH-TAMB probes failed to discover target mRNA that these cells may have produced. The exterior-facing cells might not be actively transcribing *mcr*A or the transcription level was distinctively low, therefore, they were not labeled by FISH-TAMB probes.

**Fig. 5.**
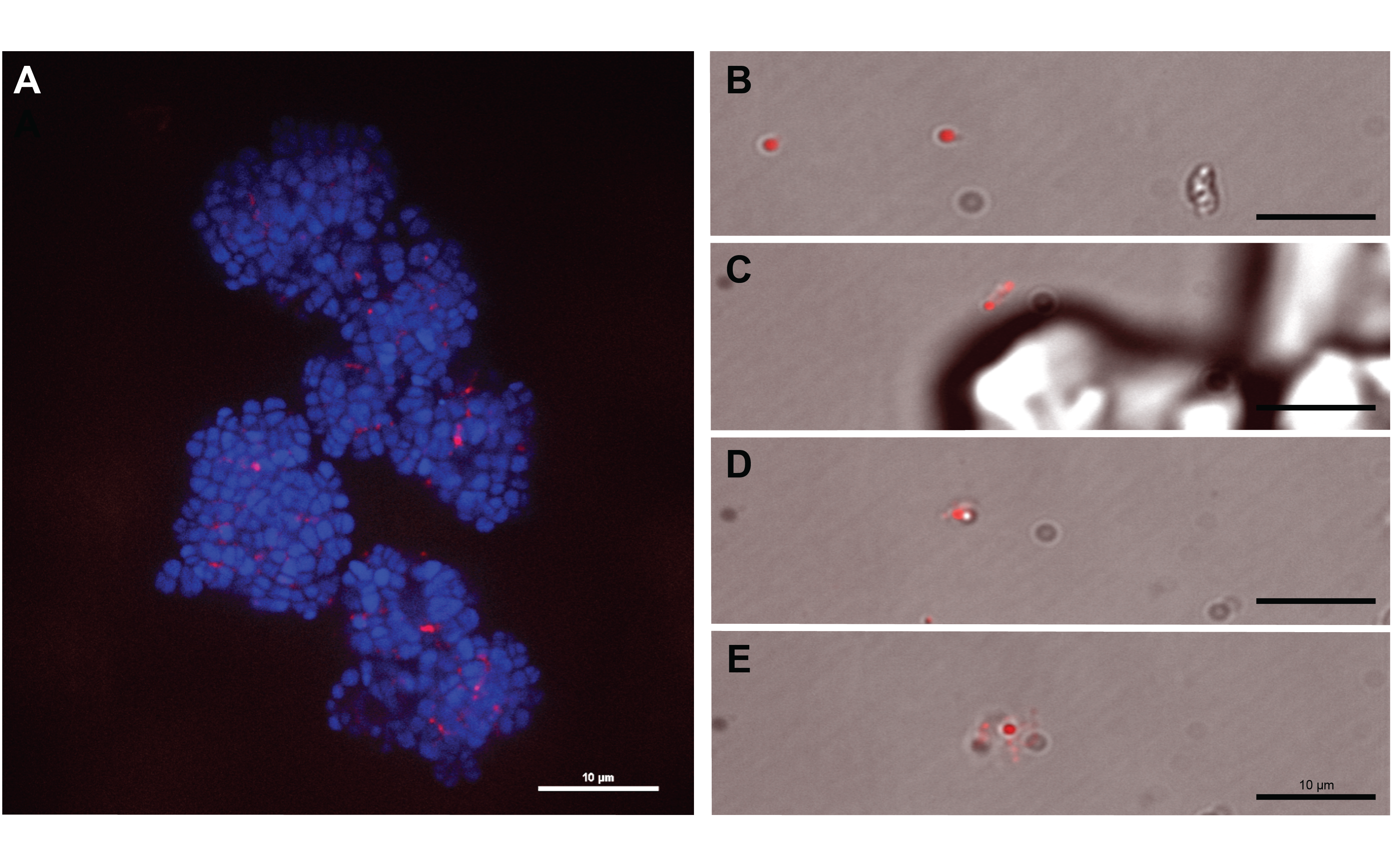
FISH-TAMB-treated methanogens labels *mcrA*-transcribing cells. Spinning disk photomicrographs of diverse surface morphologies in FISH-TAMB-treated methanogenic enrichments. **(A)** *M. barkeri* aggregate treated with 1 μM FISH-TAMB probes. Cellular autofluorescence and Cy5 fluorescence were excited with 405 nm and 647 nm lasers, respectively. Image snapped using 518 nm and 670 nm emission filters. **(B-E)** BE326 BH2-Conc methanogenic enrichments were incubated for 15 minutes with 1 μM FISH-TAMB probes and imaged every minute for 14 hours. Sample images are from the first minute of monitoring. Cy5 fluorescence was captured according to above-described parameters and integrated in real time with bright field image collection. FISH-TAMB demonstrates indiscriminate hybridization to **(B)** single cells, **(C)** physically-associated cells, **(D)** labeled and unlabeled cell pair, and **(E)** cell aggregate. Positively labeled cells indicated in red. All samples were maintained under a 100% CO_2_ atmosphere during imaging. 100x magnification. Scale bar 10 μm.

We did not assess the community composition of the BE326 BH2-Conc enrichments. However, ∼3 % of total cells were labeled by FISH-TAMB, a greater fraction than the 0.4 to 0.5% of methanogens and ANMEs reported by 16S rRNA amplicon and metagenomics data for this site (Simkus *et al.*, 2016; Lau *et al.*, 2016). Spinning disk photomicroscopy revealed FISH-TAMB-labeled cells from the BE326 BH2-Conc methanogenic enrichment exhibited small coccoidal morphologies up to 1 μm in diameter (Fig. 5 B-D), though cell aggregates were apparent in the sample (Fig. 5E). This range is consistent with cell sizes estimated from forward-scattered light area (FSC-A) distributions of reference fluorescent microspheres in flow cytometry runs (data not shown). Cy5 and bright field composite micrographs from time-lapse imaging show the presence of non-fluorescent cells in solution for the 14-hour monitoring period (Fig. 5 B-E). Fluorescence intensity of all cell morphologies was significantly reduced within 2 hours of hybridization (Fig. 6). However, single planktonic cells and cell pairs maintained discernable fluorescence for approximately 6 hours (Fig. 6 A, C).

**Fig. 6.**
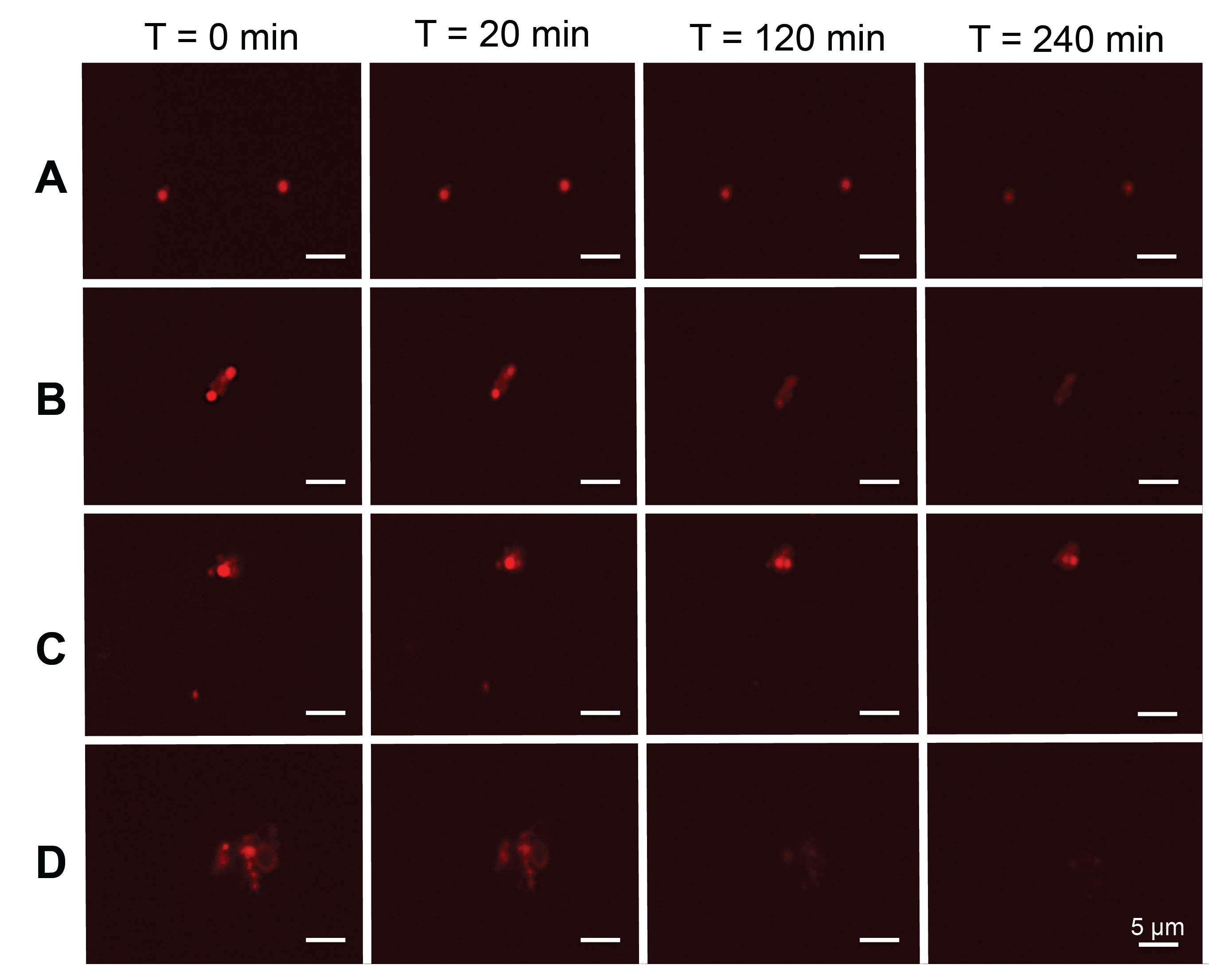
Fluorescence lifetime of Cy5 in FISH-TAMB-hybridized cells. BE326 BH2-Conc methanogenic enrichments were incubated anaerobically with 1 μM FISH-TAMB probes and subsequently imaged via spinning disk photomicroscopy. Samples were excited with a 647 nm laser line and analyzed at 670 nm under a 100% CO_2_ atmosphere. Micrographs were snapped every minute for 14 hours. Micrographs here represent the first four hours of observation. **(A)** Single cells. **(B)** Physically-associated cells. **(C)** Cell pair in which an unlabeled cell becomes labeled between 20 and 120 minutes. **(D)** Cell aggregate. 100x magnification.

FISH-TAMB-labeled cells appeared in view over the course of imaging (*SI* GIF S1), though it remains uncertain whether the appearance of these cells is due to momentary FISH-TAMB hybridization or settling of previously labeled cells into the focal plane. Weakly autofluorescent organic matter was present in aggregates. However, these emission signals were significantly weaker than that of FISH-TAMB-labeled cells (Fig. 6D).

These results demonstrated that FISH-TAMB probes enter both bacterial and archaeal cells with no harmful effect, as cells treated with FISH-TAMB remain alive. MB probes released from R9 selectively hybridize with target mRNA transcripts, if present, in the cytoplasm, emitting above-background fluorescent signals for detection.

### Cell viability of FISH-TAMB-treated cultures

Growth curves of FISH-TAMB-treated and untreated cells are presented in Fig. 7. Both FISH-TAMB-treated and untreated *E. coli* cells exhibited an extended lag phase (∼5 hours) and similar, albeit slow, growth. *E. coli* mcrA^+^ grew at μ_control_ = 0.16 h^-1^, μ_FISH-TAMB_ = 0.14 h^-1^ whereas *E. coli* pmoA^+^ grew at μ_control_ = 0.14 h^-1^, μ_FISH-TAMB_ = 0.12 h^-1^. These decreased growth rates relative to plasmid-free *E. coli* (Sezonov *et al.*, 2007; Gil-Turnes *et al.*, 2001) indicated growth inhibitory effects of the *mcr*A and *pmo*A inserts and hinted the difficulty of expressing *mcr*A in *E. coli* for FISH-TAMB detection. Doubling times for both control and FISH-TAMB-treated *M. barkeri* were ∼ 21 hours (μ = 0.03 h^-1^), which are consistent with previous reports of hydrogenotrophic *M. barkeri* growth (Maestrojuan and Boone, 1991). In both cases of *E. coli* and *M. barkeri* cells, FISH-TAMB treatments resulted in no inhibitory effects on sustained cellular viability (Fig. 7).

**Fig. 7.**
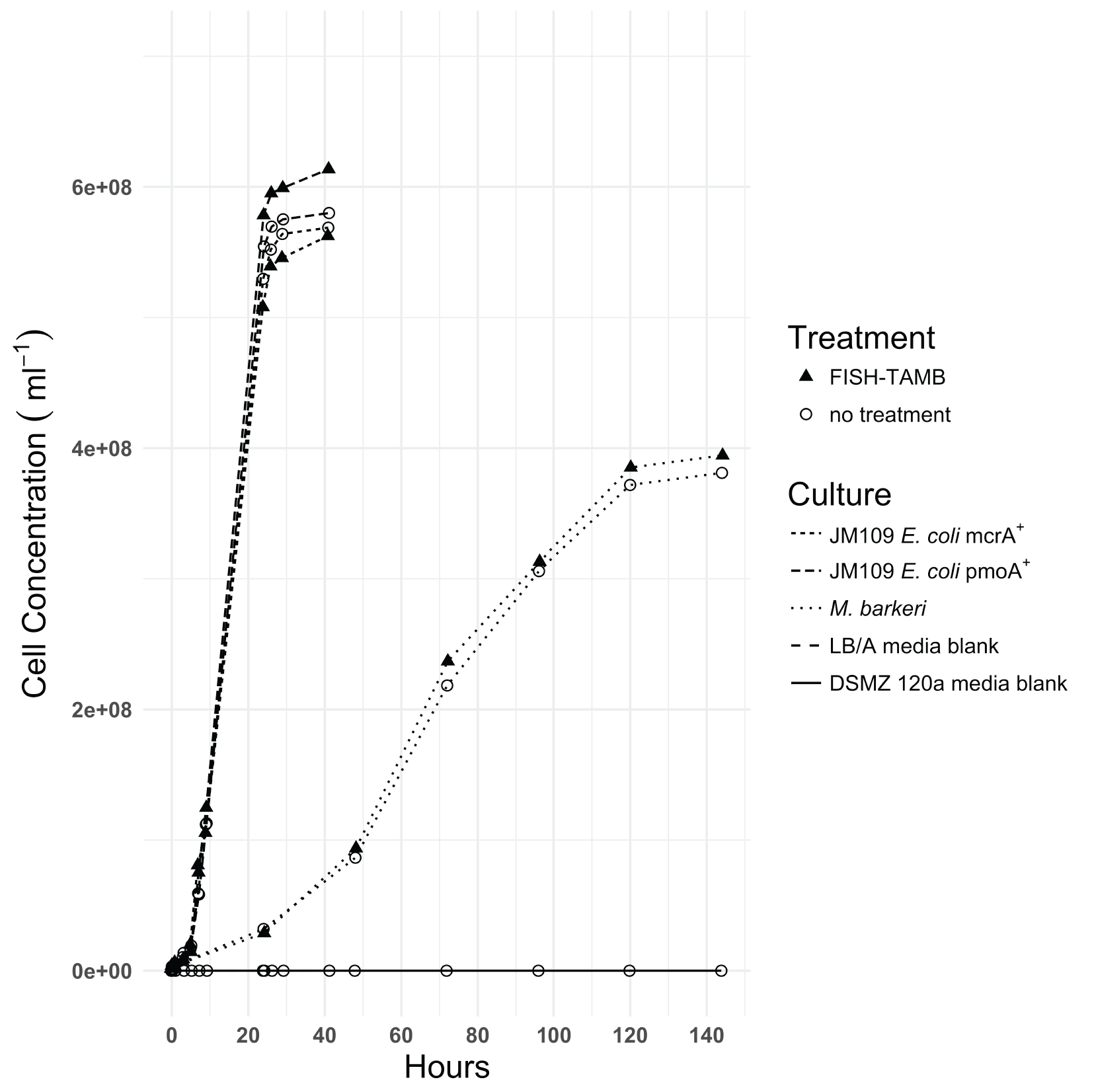
FISH-TAMB viability assessment by growth curve analysis. *E. coli* mcrA^+^, *E. coli* pmoA^+^, and *M. barkeri* cultures (∼10^6^ cells ml^-1^) were incubated with 1 μM FISH-TAMB probes and inoculated into their respective growth media. Growth was measured spectrophotometrically (OD_600_ for *E. coli*, OD_550_ for *M. barkeri*) and growth rates compared to untreated cultures.

### Implications of FISH-TAMB for microbial ecology

Cellular fixation with paraformaldehyde and ethanol is a traditional step in the FISH protocol that stabilizes cell integrity for efficient membrane permeabilization, but at the expense of DNA-protein crosslinking and potential downstream sequencing bias (Amann *et al.*, 1995; Yilmaz *et al.*, 2010). By targeting living cells, FISH-TAMB is capable of identifying cells without nucleic acid modification. In principle, FISH-TAMB can identify cells belonging active taxa from mixed microbial communities including low abundance and slow growing populations based on the expression of any target functional gene. Coupled with fluorescence-activated cell sorting, FISH-TAMB offers promising utility to isolate rare, but important, taxa for sub-cultivation and cost-efficient deep sequencing surveys.

The identification of active cells within aggregates offers opportunities to further investigate *in situ* syntrophic interactions between target cells and their physically associated partners. ANMEs and sulfate reducers are one such example of a syntrophic microbial consortium wherein the taxonomy of the implicated organisms is well described, but the mechanisms by which they perform anaerobic CH_4_ oxidation remain under debate (Valentine and Reeburgh, 2000; Hoehler *et al.*, 1994; Sørensen *et al.*, 2001; Moran *et al.*, 2008; Milucka *et al.*, 2012; McGlynn *et al.*, 2015; Wegener *et al.*, 2015). By coupling FISH-TAMB with other techniques such as Halogen In Situ Hybridization-Secondary Ion Mass Spectroscopy (HISH-SIMS) (Musat et al. 2008) and metatranscriptomics, it may be possible to improve our understanding of how myriad environmental stressors impact functional gene expression and the transfer of metabolites among cells involved in these metabolic consortia.

This study demonstrates the success of the FISH-TAMB methodology in identifying active prokaryotic cells by mRNA labeling with unnoticeable impedance to cell growth. FISH-TAMB successfully labeled *mcr*A mRNA expressed in live Bacteria and Archaea of diverse morphologies including planktonic single cells and cell aggregates. Differentiated labeling of target mRNA from cells in enrichment cultures promises applicability in utilizing FISH-TAMB to investigate active metabolic players of interest in natural microbial communities. The application to living cells suggests improved sensitivity to active minority populations that may otherwise be discriminated against detection by traditional FISH techniques. Recovery of rare taxa allows for deeper and more cost-efficient sequencing coverage of these groups in downstream “omics” applications. FISH-TAMB is an innovative step towards identifying the contributions of all microorganisms, particularly members of the rare biosphere and microbial dark matter, to biogeochemical cycling, and delineating syntrophic interactions between physically associated microorganisms.

## ACKNOWLEDGEMENTS

We are indebted to Sibanye Gold, Ltd. and the staff at the Beatrix Gold Mine for their hospitality and granting us continued access to the BE326 BH2 borehole. We would like to thank Mike Pullin, Gilbert Tetteh, Sarah Hendrickson and Olukayode Kuloyo for their field assistance, and Gilbert Tetteh’s work in isolating and amplifying *pmo*A genes from BE326 BH2. Thank you to Peter Jaffe and Melany Ruiz Uriguen for sharing access to lab equipment and to Reika Yokochi and Purtschert for offering their “Little Eddie” gas stripper for collecting gas samples at BE326 BH2.

## REFERENCES

Amann RI, Krumholz L, Stahl DA. (1990). Fluorescent-oligonucleotide probing of whole cells for determinative, phylogenetic, and environmental studies in microbiology. J Bacteriol 172: 762–770.

Amann RI, Ludwig W, Schleifer KH. (1995). Phylogenetic identification and in situ detection of individual microbial cells without cultivation. Microbiol Rev 59: 143–69.

Bao G, Rhee WJ, Tsourkas A. (2009). Fluorescent probes for live-cell RNA detection. Annu Rev Biomed Eng 11: 25–47.

Benson RC, Meyer R a, Zaruba ME, McKhann GM. (1979). Cellular autofluorescence--is it due to flavins? J Histochem Cytochem 27: 44–48.

Bryant MP, Boone DR. (1987). Emended Description of Strain MST(DSM 800T), the Type Strain of Methanosarcina barkeri. Int J Syst Bacteriol 37: 169–170.

Christensen H, Hansen M, Sørensen J. (1999). Counting and size classification of active soil bacteria by fluorescence in situ hybridization with an rRNA oligonucleotide probe. Appl Environ Microbiol 65: 1753–1761.

DeLong E, Wickham G, Pace N. (1989). Phylogenetic stains: ribosomal RNA-based probes for the identification of single cells. Science (80-) 243: 1360–1363.

Doddema HJ, Vogels GD. (1978). Improved identification of methanogenic bacteria by fluorescence microscopy. Appl Environ Microbiol 36: 752–754.

Dolfing J, Mulder JW. (1985). Comparison of methane production rate and coenzyme f(420) content of methanogenic consortia in anaerobic granular sludge. Appl Environ Microbiol 49: 1142–5.

Evans PN, Parks DH, Chadwick GL, Robbins SJ, Orphan VJ, Golding SD, et al (2015). Methane metabolism in the archaeal phylum Bathyarchaeota revealed by genome-centric metagenomics. Science 350: 434–8.

Gil-Turnes C, Conceição FR, Dellagostin OA. (2001). PRODUCTION OF PCB01, A PLASMID FOR DNA IMMUNIZATION AGAINST THE ADHESIN OF ESCHERICHIA COLI K88AB. Brazilian J Microbiol 32: 225–228.

Goel G, Kumar A, Puniya AK, Chen W, Singh K. (2005). Molecular beacon: a multitask probe. J Appl Microbiol 99: 435–442.

Golding I, Paulsson J, Zawilski SM, Cox EC. (2005). Real-time kinetics of gene activity in individual bacteria. Cell 123: 1025–1036.

Hallam SJ, Girguis PR, Preston CM, Richardson PM, DeLong EF. (2003). Identification of methyl coenzyme M reductase A (mcrA) genes associated with methane-oxidizing archaea. Appl Environ Microbiol 69: 5483–5491.

Hatzenpichler R, Connon SA, Goudeau D, Malmstrom RR, Woyke T, Orphan VJ. (2016). Visualizing in situ translational activity for identifying and sorting slow-growing archaeal-bacterial consortia. Proc Natl Acad Sci 113: E4069–E4078.

Hendrickson EL, Leigh JA. (2008). Roles of coenzyme F420-reducing hydrogenases and hydrogen-and F420-dependent methylenetetrahydromethanopterin dehydrogenases in reduction of F420 and production of hydrogen during methanogenesis. J Bacteriol 190: 4818–4821.

Hoehler TM, Alperin MJ, Albert DB, Martens S. C. (1994). Field and laboratory studies of methane oxidation in an anoxic sediment: evidence for a methanogen-sulfate-reducer consortium. Glob Biogeochem Cycles 8: 451–464.

Holmes AJ, Costello A, Lidstrom ME, Murrell JC. (1995). Evidence that participate methane monooxygenase and ammonia monooxygenase may be evolutionarily related. FEMS Microbiol Lett 132: 203–208.

Jen CJ, Chou CH, Hsu PC, Yu SJ, Chen WE, Lay JJ, et al. (2007). Flow-FISH analysis and isolation of clostridial strains in an anaerobic semi-solid bio-hydrogen producing system by hydrogenase gene target. Appl Microbiol Biotechnol 74: 1126–1134.

Kalyuzhnaya MG, Zabinsky R, Bowerman S, Baker DR, Lidstrom ME, Chistoserdova L. (2006). Fluorescence in situ hybridization-flow cytometry-cell sorting-based method for separation and enrichment of type I and type II methanotroph populations. Appl Environ Microbiol 72: 4293–4301.

Karner M, Fuhrman JA. (1997). Determination of active marine bacterioplankton: A comparison of universal 16S rRNA probes, autoradiography, and nucleoid staining. Appl Environ Microbiol 63: 1208–1213.

Larsson HM, Lee ST, Roccio M, Velluto D, Lutolf MP, Frey P, et al. (2012). Sorting Live Stem Cells Based on Sox2 mRNA Expression. PLoS One 7. e-pub ahead of print, doi:10.1371/journal.pone.0049874.

Lau MCY, Cameron C, Magnabosco C, Brown CT, Schilkey F, Grim S, et al. (2014). Phylogeny and phylogeography of functional genes shared among seven terrestrial subsurface metagenomes reveal N-cycling and microbial evolutionary relationships. Front Microbiol 5. e-pub ahead of print, doi:10.3389/fmicb.2014.00531.

Lau MCY, Kieft TL, Kuloyo O, Linage-Alvarez B, van Heerden E, Lindsay MR, et al. (2016). An oligotrophic deep-subsurface community dependent on syntrophy is dominated by sulfur-driven autotrophic denitrifiers. Proc Natl Acad Sci U S A 201612244.

Lazar CS, Baker BJ, Seitz KW, Teske AP. (2017). Genomic reconstruction of multiple lineages of uncultured benthic archaea suggests distinct biogeochemical roles and ecological niches. ISME J. http://dx.doi.org/10.1038/ismej.2016.189.

Liu BR, Huang Y-W, Lee H-J. (2013a). Mechanistic studies of intracellular delivery of proteins by cell-penetrating peptides in cyanobacteria. BMC Microbiol 13: 57.

Liu BR, Liou JS, Huang YW, Aronstam RS, Lee HJ. (2013b). Intracellular Delivery of Nanoparticles and DNAs by IR9 Cell-penetrating Peptides. PLoS One 8. e-pub ahead of print, doi:10.1371/journal.pone.0064205.

Lueders T, Chin KJ, Conrad R, Friedrich M. (2001). Molecular analyses of methyl-coenzyme M reductase α-subunit (mcrA) genes in rice field soil and enrichment cultures reveal the methanogenic phenotype of a novel archaeal lineage. Environ Microbiol 3: 194–204.

Luesken FA, Zhu B, van Alen TA, Butler MK, Diaz MR, Song B, et al. (2011). pmoA primers for detection of anaerobic methanotrophs. Appl Environ Microbiol 77: 3877–3880.

Luton PE, Wayne JM, Sharp RJ, Riley PW. (2002). The mcrA gene as an alternative to 16S rRNA in the phylogenetic analysis of methanogen populations in landfill. Microbiology 148: 3521–3530.

Maestrojuan GM, Boone DR. (1991). Characterization of Methanosarcina barkeri MST and 227, Methanosarcina mazei S-6T, and Methanosarcina vacuolata Z-761T. Int J Syst Bacteriol 41: 267–274.

Magnabosco C, Tekere M, Lau MCY, Linage B, Kuloyo O, Erasmus M, et al. (2014). Comparisons of the composition and biogeographic distribution of the bacterial communities occupying South African thermal springs with those inhabiting deep subsurface fracture water. Front Microbiol 5. e-pub ahead of print, doi:10.3389/fmicb.2014.00679.

McGlynn SE, Chadwick GL, Kempes CP, Orphan VJ. (2015). Single cell activity reveals direct electron transfer in methanotrophic consortia. Nature 526: 531–535.

Milucka J, Ferdelman TG, Polerecky L, Franzke D, Wegener G, Schmid M, et al. (2012). Zero valent sulphur is a key intermediate in marine methane oxidation. Nature 491: 541–546.

Moran JJ, Beal EJ, Vrentas JM, Orphan VJ, Freeman KH, House CH. (2008). Methyl sulfides as intermediates in the anaerobic oxidation of methane. Environ Microbiol 10: 162–173.

Mota CR, So MJ, de los Reyes FL. (2012). Identification of Nitrite-Reducing Bacteria Using Sequential mRNA Fluorescence In Situ Hybridization and Fluorescence-Assisted Cell Sorting. Microb Ecol 64: 256–267.

Nitin N, Santangelo PJ, Kim G, Nie S, Bao G. (2004). Peptide-linked molecular beacons for efficient delivery and rapid mRNA detection in living cells. Nucleic Acids Res 32: e58.

Pernthaler A, Amann R. (2004). Simultaneous Fluorescence In Situ Hybridization of mRNA and rRNA in Environmental Bacteria Simultaneous Fluorescence In Situ Hybridization of mRNA and rRNA in Environmental Bacteria. Appl Environ Microbiol 70: 5426–5433.

Pernthaler A, Preston CM, Pernthaler J, DeLong EF, Amann R. (2002). Comparison of fluorescently labeled oligonucleotide and polynucleotide probes for the detection of pelagic marine bacteria and archaea. Appl Environ Microbiol 68: 661–667.

Renggli S, Keck W, Jenal U, Ritz D. (2013). Role of Autofluorescence in Flow Cytometric Analysis of Escherichia coli Treated with Bactericidal Antibiotics. J Bacteriol 195: 4067–4073.

Rinke C, Schwientek P, Sczyrba A, Ivanova NN, Anderson IJ, Cheng J-F, et al. (2013). Insights into the phylogeny and coding potential of microbial dark matter. Nature 499: 431–437.

Saïda F, Uzan M, Odaert B, Bontems F. (2006). Expression of highly toxic genes in E. coli: special strategies and genetic tools. Curr Protein Pept Sci 7: 47–56.

Santangelo P, Nitin N, Bao G. (2006). Nanostructured probes for RNA detection in living cells. Ann Biomed Eng 34: 39–50.

Seitz KW, Lazar CS, Hinrichs K-U, Teske AP, Baker BJ. (2016). Genomic reconstruction of a novel, deeply branched sediment archaeal phylum with pathways for acetogenesis and sulfur reduction. ISME J 10: 1696–1705.

Sezonov G, Joseleau-Petit D, D’Ari R. (2007). Escherichia coli physiology in Luria-Bertani broth. J Bacteriol 189: 8746–8749.

Simkus DN, Slater GF, Lollar BS, Wilkie K, Kieft TL, Magnabosco C, et al. (2016). Variations in microbial carbon sources and cycling in the deep continental subsurface. Geochim Cosmochim Acta 173: 264–283.

Sogin ML, Sogin ML, Morrison HG, Morrison HG, Huber J a, Huber J a, et al. (2006). Microbial diversity in the deep sea and the underexplored ‘rare biosphere’. Proc Natl Acad Sci U S A 103: 12115–20.

Sokol DL, Zhang X, Lu P, Gewirtz a M. (1998). Real time detection of DNA.RNA hybridization in living cells. Proc Natl Acad Sci U S A 95: 11538–43.

Sørensen KB, Finster K, Ramsing NB. (2001). Thermodynamic and Kinetic Requirements in Anaerobic Methane Oxidizing Consortia Exclude Hydrogen, Acetate, and Methanol as Possible Electron Shuttles. Microb Ecol 42: 1–10.

Spang A, Saw JH, Jørgensen SL, Zaremba-Niedzwiedzka K, Martijn J, Lind AE, et al. (2015). Complex archaea that bridge the gap between prokaryotes and eukaryotes. Nature 521: 173–179.

Steinberg LM, Regan JM. (2008). Phylogenetic comparison of the methanogenic communities from an acidic, oligotrophic fen and an anaerobic digester treating municipal wastewater sludge. Appl Environ Microbiol 74: 6663–71.

Tzschaschel BD, Guzmán CA, Timmis KN, Lorenzo V de. (1996). An Escherichia coli hemolysin transport system-based vector for the export of polypeptides: Export of shiga like toxin IIeB subunit by Salmonella typhimurium aroA. Nat Biotechnol 14: 765–769.

Valentine DL, Reeburgh WS. (2000). New perspectives on anaerobic methane oxidation. Env Microbiol 2: 477–484.

Vanwonterghem I, Evans PN, Parks DH, Jensen PD, Woodcroft BJ, Hugenholtz P, et al. (2016). Methylotrophic methanogenesis discovered in the archaeal phylum Verstraetearchaeota. Nat Microbiol 1: 16170.

Wegener G, Krukenberg V, Riedel D, Tegetmeyer HE, Boetius A. (2015). Intercellular wiring enables electron transfer between methanotrophic archaea and bacteria. Nature 526: 587–590.

Williams SC, Hong Y, Danavall DCA, Howard-Jones MH, Gibson D, Frischer ME, et al. (1998). Distinguishing between living and nonliving bacteria: Evaluation of the vital stain propidium iodide and its combined use with molecular probes in aquatic samples. J Microbiol Methods 32: 225–236.

Yilmaz S, Haroon MF, Rabkin BA, Tyson GW, Hugenholtz P. (2010). Fixation-free fluorescence in situ hybridization for targeted enrichment of microbial populations. ISME J 4: 1352–1356.

Zaremba-Niedzwiedzka K, Caceres EF, Saw JH, Bäckström D, Juzokaite L, Vancaester E, et al. (2017). Asgard archaea illuminate the origin of eukaryotic cellular complexity. Nature 541: 353–358.

